# Tianeptine, but not fluoxetine, decreases avoidant behavior in a mouse model of early developmental exposure to fluoxetine

**DOI:** 10.1101/2021.01.19.427351

**Authors:** Elizabeth A Pekarskaya, Emma S Holt, Jay A Gingrich, Mark S Ansorge, Jonathan A Javitch, Sarah E Canetta

**Author notes:** denotes co-corresponding authorship. denotes co-first authorship (Elizabeth A Pekarskaya, Emma S Holt).

## Abstract

Depression and anxiety are two of the most common mental health disorders, often sharing symptoms and administrations. Most pharmacological agents available to treat these disorders target monoamine systems. Currently, finding the most effective treatment for an individual is a process of trial and error. Therefore, to better understand how disease etiology may predict treatment response, we studied mice exposed *developmentally* to the selective serotonin reuptake inhibitor (SSRI) fluoxetine (FLX). These mice show the murine equivalent of anxiety- and depression-like symptoms in adulthood and here we report that these mice are also behaviorally resistant to the antidepressant-like effects of adult SSRI administration. We investigated whether tianeptine (TIA), which exerts its therapeutic effects through the mu-opioid receptor (MOR) instead of directly targeting monoaminergic systems, would be more effective in this model.

We injected C57BL/6J (C57) pups with either FLX (10 mg/kg, i.p) or vehicle from postnatal (PN) day 2 to 11, a period in which mouse brain development parallels that of the third trimester of a human pregnancy. Prior work established that adult 129SvEv (129) mice exposed to FLX in this time period (PN-FLX) showed increased avoidant and decreased hedonic behaviors, which correspond to anxiety- and depressive-like symptoms in humans, respectively. We performed baseline testing in adulthood in C57 PN-FLX animals and confirmed a similar avoidant phenotype to that reported in 129 PN-FLX mice. We then treated these animals with chronic FLX (18 mg/kg in the drinking water) and evaluated effects on two tasks that measure avoidant behavior – the open field and novelty suppressed feeding (NSF) tasks. This administration failed to improve, and even exacerbated, avoidance symptoms in PN-FLX mice. The same animals then underwent chronic administration with TIA (30 mg/kg, 2x/day, i.p.) as an alternative treatment strategy. TIA administration decreased avoidance behavior as measured in the open field and NSF. Overall, this demonstrates that TIA may be a promising alternative treatment to typical antidepressants, especially in patients whose serotonergic system has been altered.

## INTRODUCTION

Depression and anxiety-related disorders are the two most common mental health conditions, and in the top ten of all health conditions contributing to disability globally^1^. This situation has been exacerbated by the recent COVID-19 pandemic, with thirty-five percent of adults currently reporting symptoms of anxiety and/or depression and rates as high as forty-six percent in young adults ages 18-29 in the US^2^. The etiology of these disorders is highly complex. Genetics, early life adversity, acute stress, traumatic events, hormones, and other factors have all been shown to increase risk^3^. While depression and anxiety-related disorders are distinguished diagnostically, they share considerable overlap in symptomatology, including anhedonia and avoidant behaviors in patients^4,5^. In addition, both conditions are highly comorbid, implying some shared neurobiological substrates.

Treatments for these conditions also often overlap, with nearly all focused on manipulating monoaminergic systems such as serotonin, norepinephrine, and dopamine. Frontline pharmacological treatments for both depression and anxiety disorders are selective serotonin reuptake inhibitors (SSRIs) such as fluoxetine (FLX)^4^. SSRIs function by blocking the serotonin transporter, thereby increasing the amount of available extracellular serotonin. Over a period of weeks, the resulting increase in serotonin is thought to help normalize mood, feelings of worthlessness, sleep disruption, and anxiety in responsive patients^6^. Unfortunately, 53% of patients show non-response and 67% show non-remission following treatment with frontline SSRIs^7^, with some experiencing a worsening of symptoms and an increased risk of suicide. Part of this administration resistance may be attributable to our current inability to identify patients with differing underlying neurobiology for whom treatments with a different pharmacological target might be more effective.

Studies in mice can help us better understand how disease etiology may predict treatment response. In mice, avoidant, anhedonic, motivated and despair-related behavioral measures are used as a proxy for anxiety- and depression-like behaviors in humans^8^. Early postnatal exposure (postnatal day 2-11) to FLX paradoxically results in increased avoidance behavior and decreased novelty-induced exploration and hedonic drive, as well as impairments in cognitive tasks, and diminished fear extinction^9^. These behavioral changes in postnatal fluoxetine-exposed (PN-FLX) mice are believed to result, at least in part, from persistent alterations in their serotonergic system^9–11^. Intriguingly, in the current study, we find that these avoidant symptoms in PN-FLX mice are not responsive to subsequent SSRI administration in adulthood.

Importantly, this early sensitive time period of SSRI exposure in mice is believed to correspond to human exposure during the third trimester *in utero*, suggesting these findings may have implications for the children of those who take SSRIs while pregnant. Indeed, a recent longitudinal cohort study in Finland found that rates of depression in the children of women who took antidepressants during pregnancy were significantly elevated compared to the children of women who had depression but never took an antidepressant or discontinued its use during pregnancy^12^. However, recommendations for the treatment of anxiety and depression in pregnant patients remain complex as this risk must be balanced with concerns about the consequences of undertreated maternal depression on the health of the patient as well as prenatal and infant care^13,14^. Given the prevalence of anxiety and depression, and the widespread usage of SSRIs as a treatment for these conditions, it is of substantial clinical relevance to understand if prenatal exposure to SSRIs might be an important aspect of medical history that would predict treatment response. In particular, one hypothesis is that this patient subpopulation may benefit from antidepressants that do not directly target serotonergic neurons via autoreceptors or transporters^15,16^.

In preclinical studies, the atypical antidepressant tianeptine (TIA) was recently shown to exert its effects on affective behaviors via activation of the mu-opioid receptor (MOR)^17,18^. Although MOR activation is responsible for both the analgesic and addictive effects of opioid pain relievers such as morphine, TIA is not euphoric at clinical doses^19^, and has been considered as a viable alternative to frontline pharmacological options that directly target monoaminergic systems for treatment-resistant depression^20^.

Based on the different hypothesized mechanisms of action between TIA and SSRIs, the goal of this study was to investigate the effectiveness of TIA at relieving avoidant behaviors associated with developmental exposure to FLX in mice. To test this, we injected C57BL/6J (C57) pups with FLX from postnatal day 2 to postnatal day 11 (P2-P11), and then administered FLX or TIA chronically in adulthood, and evaluated the efficacy of each compound on normalizing avoidant behavior in the open field and novelty suppressed feeding task. Overall, our study is designed to assess and compare efficacy of FLX versus TIA in a mouse model of gestational SSRI exposure, which may have broader implications for affective disorder subtypes with a shared etiology that have been resistant to SSRI treatment.

## METHODS

### Subjects

Male and female C57BL/6J (C57) mice (Jackson Laboratories, Bar Harbor, ME) were used to breed a total of 32 male and 38 female pups. Prior to giving birth, dams were individually housed. Pups within each litter were divided into PN-FLX and PN-VEH groups. The PN-FLX group consisted of 18 males and 21 females (39 in total), and the PN-VEH group consisted of 14 males and 17 females (31 total). Pups were weaned at postnatal day 21 with PN-FLX and PN-VEH mice in each cage. After postnatal day 90, cages were pseudo-randomly sorted into chronic adulthood administration groups, first FLX or VEH, and then subsequently for TIA or VEH. For the FLX administration experiments, 20 PN-FLX (9 males, 11 females) and 16 PN-VEH (7 males, 9 females) mice received FLX, and 19 PN-FLX (9 males, 10 females) and 15 PN-VEH (7 males, 8 females) mice received the VEH. Sexes were combined for analysis. For the TIA administration experiments, 20 PN-FLX (10 males, 10 females) and 16 PN-VEH (8 males, 8 females) received TIA, and 19 PN-FLX (8 males, 11 females) and 15 PN-VEH (6 males, 9 females) received VEH. 129SvEv (129) mice were purchased from Taconic (Taconic Farms, Germantown, New York, USA). Of 57 PN-FLX mice, 30 received FLX as adults and 27 received VEH (sexes combined). Of 68 PN-VEH mice, 32 received FLX and 36 received VEH. 17 animals did not receive any postnatal administration and are referred to as naïve. Of these mice, 8 received FLX and 9 received the VEH. The 129 mice did not receive TIA administration following the chronic FLX administration and were only used to compare chronic adult FLX administration versus VEH. Mice were group housed 3-5 to a cage, were kept on a 12-hour light/dark cycle, and unless otherwise noted, given *ad libitum* food and water. All behavioral assays were performed during the light portion of the cycle. All procedures were carried out in accordance with guidelines approved by the Institutional Animal Care and Use Committees at Columbia University and the New York State Psychiatric Institute.

### Drug administrations

#### Postnatal fluoxetine

From P2-P11, pups were given either 10 mg/kg (2 mg/ml) Fluoxetine HCL (Anawa, Kloten, Switzerland) or a 0.9% sterile saline vehicle through intraperitoneal injection once per day. The 2 mg/ml solution for injection was prepared daily by diluting a 4 mg/ml sterile stock solution of fluoxetine in milliQ water stored at 4°C with 1.8% sterile saline. The drug was administered at 5 μl/g weight of pup.

#### Adulthood fluoxetine

Cages with co-housed PN-FLX and PN-VEH mice were pseudorandomly organized into adult administration groups. Fluoxetine HCL was administered through the drinking water for three weeks once all the animals had matured beyond P90. Behavioral testing began after three weeks of administration, and FLX administration continued during testing. The 18 mg/kg solution appropriate for C57 mice^21,22^ was prepared by dissolving FLX into tap water. A 10 mg/kg solution was used for 129 mice^23^. The respective vehicle was the tap water alone. Opaque water bottles were used to reduce FLX light exposure. Bottles were changed and weighed every few days to track consumption.

#### Tianeptine

A two-week washout period followed completion of post-FLX behavioral testing before beginning TIA administration. Cages were pseudorandomly reorganized into TIA and VEH groups, ensuring distribution across both PN-FLX and PN-VEH groups, and those which had received FLX or VEH during the adult administration period. There was no effect of prior adulthood FLX administration on behavior following TIA administration. A 30 mg/kg solution of tianeptine sodium salt (Qingdao Sigma Chemical Co., purity verified by NMR analysis) was delivered at a dose of 0.1 ml/10 g mouse weight via intraperitoneal injection twice per day. Tianeptine sodium salt was dissolved in 0.9% sterile saline and stored at room temperature for up to three days. The respective vehicle was 0.9% sterile saline.

### Behavior

#### Open field test (OFT)

Mice were placed in 42 × 42 × 38 cm plexiglass enclosures for 60 minutes. Activity was measured using MotorMonitor software (Kinder Scientific, Poway, CA), which recognizes infrared beam breaks in the enclosure. Specifically, the number of ambulatory movements, the number of rearing instances, and time spent in the center were analyzed. The center was defined as a centered 21 × 21 cm grid. Each of these variables was summed over the first 10 minutes or total 60 minutes.

#### Forced swim test (FST)

Mice were habituated to the experimental room for 1 hour prior to testing. Mice swam for 6 minutes in containers filled with 2000 ml of 25 - 30°C tap water. Temperature of the water was monitored and the water was changed as needed. Video was recorded and the first 2 minutes were analyzed for immobility using Videotrack software (ViewPoint, France).

#### Novelty suppressed feeding (NSF)

Mice were food deprived 15 hours prior to the beginning of their NSF test. The detailed protocol and description of the containers used can be found in Samuels & Hen, 2011^24^. A pellet of food was secured onto a white surface that was placed in the midst of familiar bedding in a novel arena under 1200 lux overhead lighting. C57 mice were allowed up to 6 minutes, 10 minutes for 129 mice, to explore the arena and bite the food pellet before timing out. Latency to bite the pellet, referred to as latency to feed, was recorded by the experimenter. This was followed by 5 minutes in the home cage, during which mice were presented with a food pellet. Both home cage latency to feed and amount of food consumed were recorded by the experimenter.

### Analysis

Unless otherwise noted, data were analyzed using Prism (GraphPad, San Diego, CA) or using custom scripts in Matlab (Mathworks, Natick, MA). Data from software that monitored activity during behavioral tests were imported into these programs to be analyzed by experimental group. Sexes were combined for statistical analysis unless a two-way ANOVA revealed a statistical interaction of sex and administration for a main finding.

## RESULTS

### C57 mice exposed to fluoxetine from P2-11 exhibit increased avoidant behavior in adulthood

Prior studies established that 129 mice exposed to FLX from P2-P11 display increased avoidant behaviors and decreased hedonic behaviors as adults^9^. A similar increase in avoidant behaviors had been demonstrated in C57 mice, treated for a slightly longer window from P4-P21^25^. The C57 strain was chosen for the current experiments because of the established efficacy of TIA for normalizing similar types of affective symptoms in a chronic corticosterone administration model^18^. Both male and female mice received either FLX or VEH injections once a day from P2-P11 (Fig. 1a; shown in purple), a developmental window in mice in which brain development is comparable to that during the third trimester in humans^9^. While there was no discernable difference between groups as pups, mature mice that had received FLX as pups (PN-FLX) weighed significantly less than their VEH counterparts (PN-VEH) (Table 1; two-way ANOVA, F(1, 66) = 0.11, P = 0.75 for interaction of sex and administration, F(1, 66) = 450.4, ****P < 0.0001 for main effect of sex, F(1, 66) = 15.59, ***P = 0.0002 for main effect of administration; Bonferroni post hoc males, *P = 0.02, n = 14 PN-VEH, 18 PN-FLX; females, *P = 0.01, n = 17 PN-VEH, 21 PN-FLX). Once these mice entered adulthood, a number of behavioral tests were performed to determine whether they had a baseline affective phenotype prior to adult antidepressant administration (Fig. 1A; shown in yellow).

**Figure 1.**
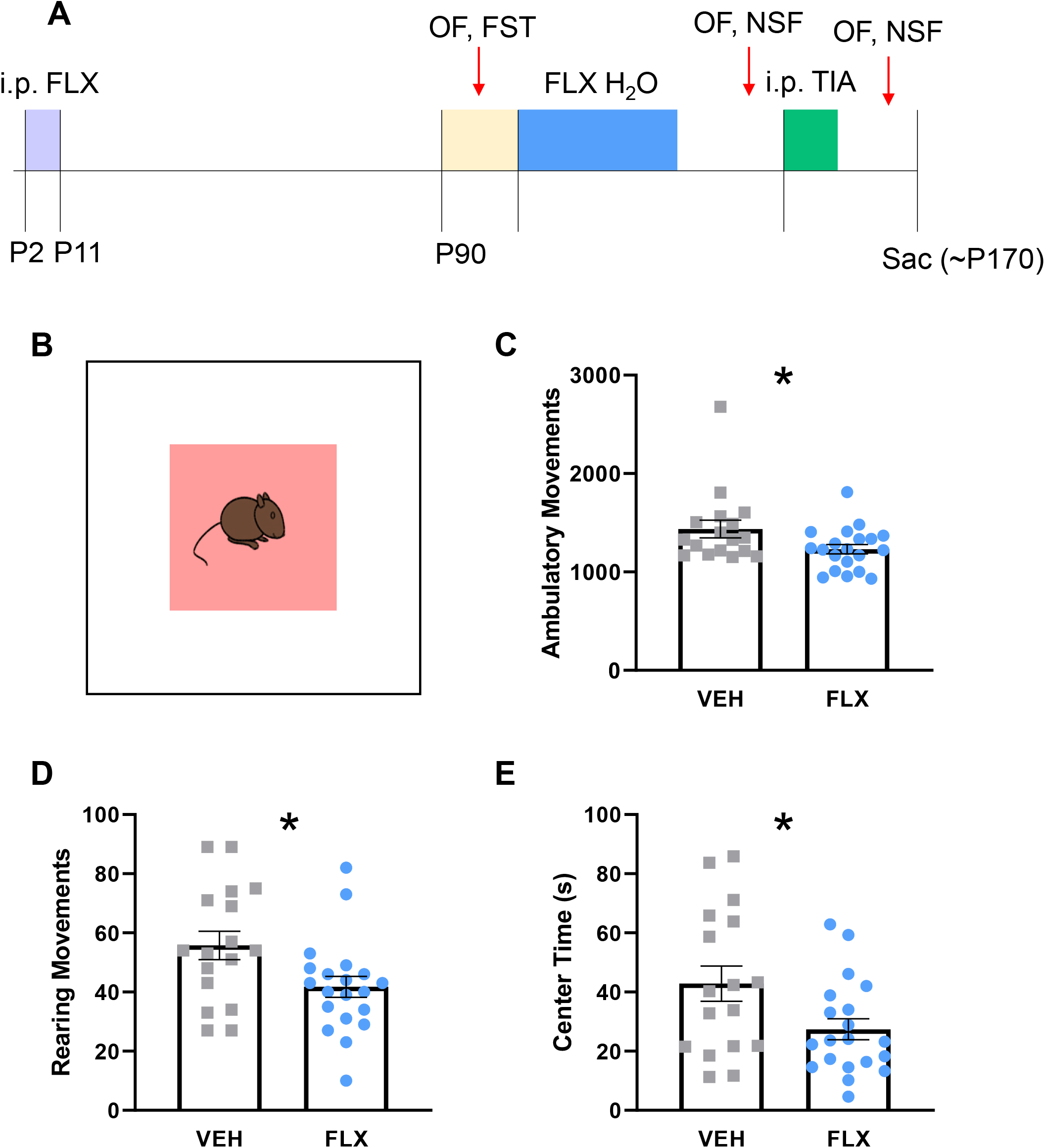
Decreased novelty-induced exploration and increased avoidance behavior in the first 10 minutes of open field following chronic adult fluoxetine treatment in post-natal fluoxetine mice. **A.** Timeline of experiment. Blue shows the period during which chronic fluoxetine was administered after which NSF and OF tests were performed. OF (open field), FST (forced swim test), NSF (novelty suppressed feeding). **B.** Diagram of open field enclosure, with center shown in red. **C.** Ambulatory movements in first 10 minutes of open field. Unpaired, two-tailed, t-test, p=0.0440. **D.** Same as C, except rearing movements are shown, unpaired two-tailed, t-test, p=0.0224. **E.** Same as C, except time spent in center is shown, unpaired, two-tailed, t-test, p=0.0268. Gray squares represent adult vehicle-treated PN FLX animals, N=17. Blue circles represent adult fluoxetine-treated PN FLX animals, N=20. Bars represent mean. Error bars represent standard error. *p<0.05.

**Table 1.**
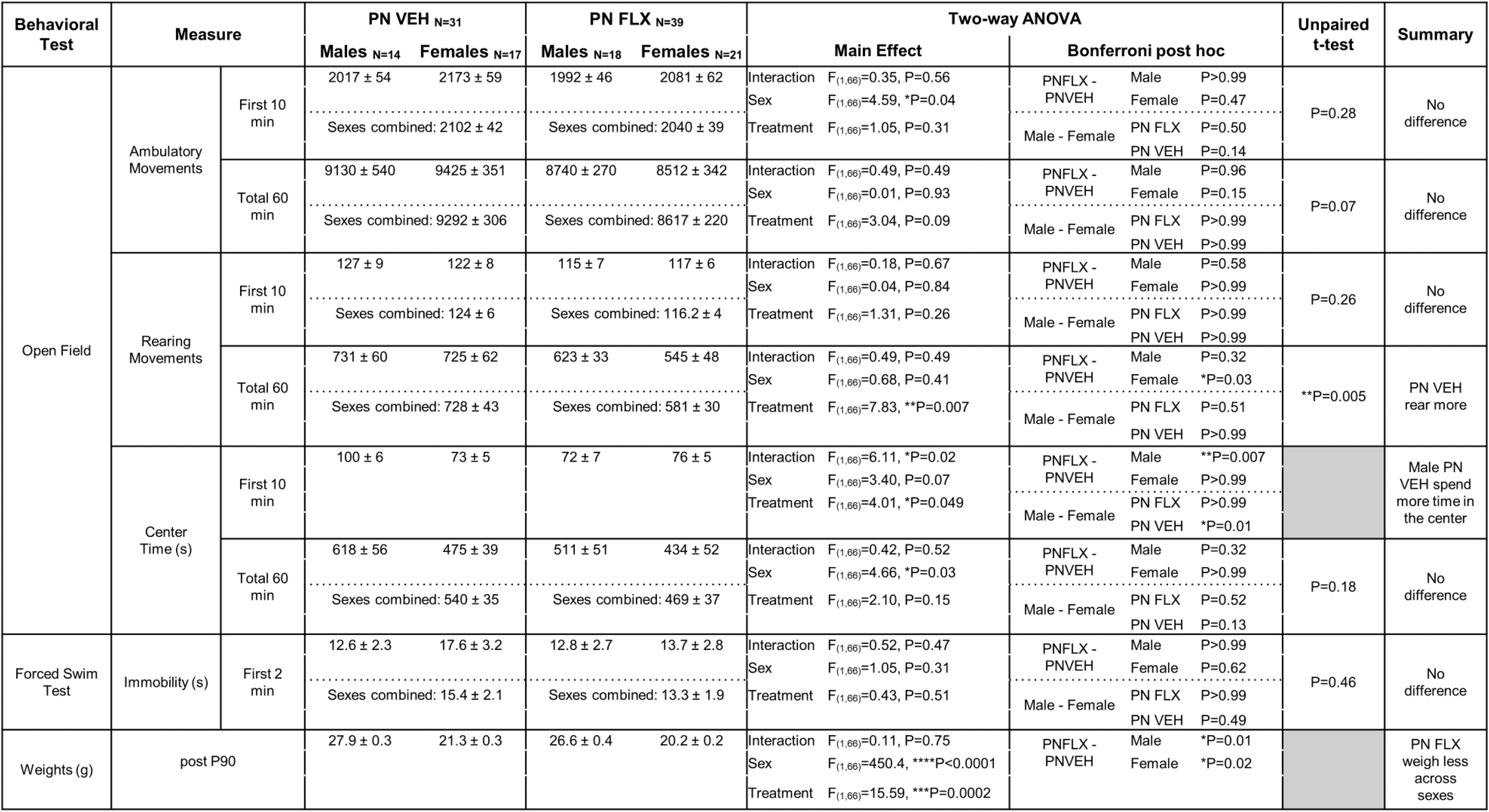
Post-natal fluoxetine mice exhibit increased avoidant behavior in adulthood. Post-natal fluoxetine (PN FLX) and post-natal vehicle (PN VEH) treated adults were tested in the OF (open field) and FST (forced swim test) for avoidant and depressive-like behaviors, respectively, following post-natal day 90. Sexes were combined for analysis where there was no statistically significant interaction between sex and postnatal treatment. PN VEH N=31, males N=14, females N=17. PN FLX N=39, males N=18, females N=21.

In the OFT, rearing movements were analyzed to measure novelty-induced exploration^26^, and time spent in the center was analyzed to measure avoidant behavior. We separately assessed these behaviors in the first 10 minutes of the task, the time period most sensitive to changes in avoidance and exploratory drive^23^, as well as the full 60 minutes. In the first 10 minutes, PN-VEH mice did not rear more than PN-FLX mice (Table 1; unpaired t-test, P=0.26, n=31 PN-VEH, 39 PN-FLX, sexes combined). However, when rearing movements were summed over the total 60 minutes, PN-VEH mice reared significantly more than PN-FLX mice (Table 1; unpaired t-test, **P=0.005, n=31 PN-VEH, 39 P-FLX, sexes combined), consistent with what was previously observed in 129 mice. This suggests that postnatal FLX exposure reduced novelty-induced exploration in C57 mice. Consistently, male PN-FLX mice spent less time in the center during the first 10 minutes compared to their PN-VEH counterparts (Table 1; two-way ANOVA, F(1, 66)=6.11, *P=0.02 for interaction of sex and drug, F(1, 66) = 3.40, P = 0.07 for main effect of sex, F(1, 66) = 4.01, *P = 0.049 for main effect of drug; Bonferroni post hoc males, **P = 0.007, n = 14 PN-VEH, 18 PN-FLX; females, P > 0.99, n = 17 PN-VEH, 21 PN-FLX). Total ambulatory movements in the OF were measured across the entire 60 minutes to asses overall locomotion and ensure that novelty induced changes in behavior in the first 10 minutes of the test were not the result of broader locomotor dysfunction. PN-FLX and PN-VEH mice did not differ in number of ambulatory movements (Table 1; first 10 minutes, unpaired t-test, P = 0.28, n = 31 PN-VEH, 39 PN-FLX, sexes combined; total 60 minutes, unpaired t-test, P = 0.07, n = 31 PN-VEH, 39 PN-FLX, sexes combined). As overall ambulation did not differ across the groups (Table 1), and a decrease in center-time during the first 10 minutes was observed, this suggests that postnatal FLX exposure resulted in an increase in avoidant behavior in male C57s.

The FST measures behavioral despair and is generally considered a proxy for depressive-like behavior in mice^27^. In the FST, the mouse is placed in a beaker of water from which it cannot escape, and the amount of time it spends struggling versus giving up and remaining immobile is used to assess despair response in the face of stress. Typically, a high percentage of time spent immobile is considered an indicator of depressive-like behavior^27,28^. PN-FLX and PN-VEH mice performed similarly in this task. Time spent immobile was equivalent between PN-FLX mice and PN-VEH mice (Table 1; unpaired t-test, P=0.46, n=31 PN-VEH, 39 PN-FLX, sexes combined). The results of the FST suggest that postnatal FLX did not produce a depressive phenotype in C57 mice. This contrasts with what has been observed in 129 mice^9^ but similar to what has been seen in C57 mice following P4-P21 PN FLX administration^25^.

Overall, early postnatal FLX exposure in C57 mice produces a persistent behavioral phenotype in adults consistent with decreased novelty-induced exploration and increased avoidance behavior, which is not due to gross abnormalities in locomotor ability.

### Chronic fluoxetine exacerbates avoidant behavior in adult mice postnatally exposed to fluoxetine

After a baseline phenotype was established, we chronically administered PN-FLX and PN-VEH mice with FLX or VEH via drinking water in the home cage for 3 weeks (Fig. 1a; shown in blue). Prior work has established this method of FLX administration as effective in normalizing avoidant behaviors in a variety of tasks including the OFT and NSF in a corticosterone-induced model of anxiety and depression-like behavior in C57 mice^22^. Following FLX administration, mice were retested in the OFT. We found that chronic FLX administration did not decrease avoidant behaviors, and instead exacerbated them. PN-FLX mice administered with FLX showed a reduction in rearing (Fig. 1d; unpaired t-test, *P = 0.02, n = 17 VEH, 20 FLX, sexes combined) and time spent exploring the center in the first 10 minutes of the OFT (Fig. 1e; unpaired t-test, *P = 0.03, n = 17 VEH, 20 FLX, sexes combined). As the effect of chronic FLX was not seen in these measures when summing across 60 minutes in the OFT (Fig. S1), we believe this reduction in exploratory behavior and increase in avoidant behavior is not due to an overall decrease in locomotor function^29^. Chronic FLX administration in PN-VEH mice led to an overall decrease in ambulation, but did not lead to a significant difference in center time in the first time 10 minutes (Fig. S2), consistent with prior literature demonstrating that chronic FLX administration leads to decreased locomotion and is not anxiolytic in unstressed C57 mice^23,30^.

Additionally, following chronic FLX or VEH administration, we tested the PN-FLX and PN-VEH mice in another test of avoidant behavior, the NSF. Chronic FLX administration has previously been shown to decrease latency to feed in the NSF in a model of corticosterone-induced anxiety- and depression-like behavior in C57 mice^22^. In contrast, chronic FLX had no effect on latency to feed in PN-FLX mice (Fig. 2b; unpaired t-test, P = 0.70, n = 17 VEH, 20 FLX, sexes combined). This was not due to insufficient hunger, as chronic FLX did not affect the amount of food consumed by PN-FLX mice in the home cage (Fig. 2c; unpaired t-test, P = 0.37, n = 17 VEH, 20 FLX, sexes combined), nor their latency to eat in the home cage (Fig. 2d; unpaired t-test, P = 0.20, n = 17 VEH, 20 FLX, sexes combined). Similarly, chronic FLX administration had no effect on PN-VEH mice compared to their VEH counterparts in the NSF (Fig. S3).

**Figure 2.**
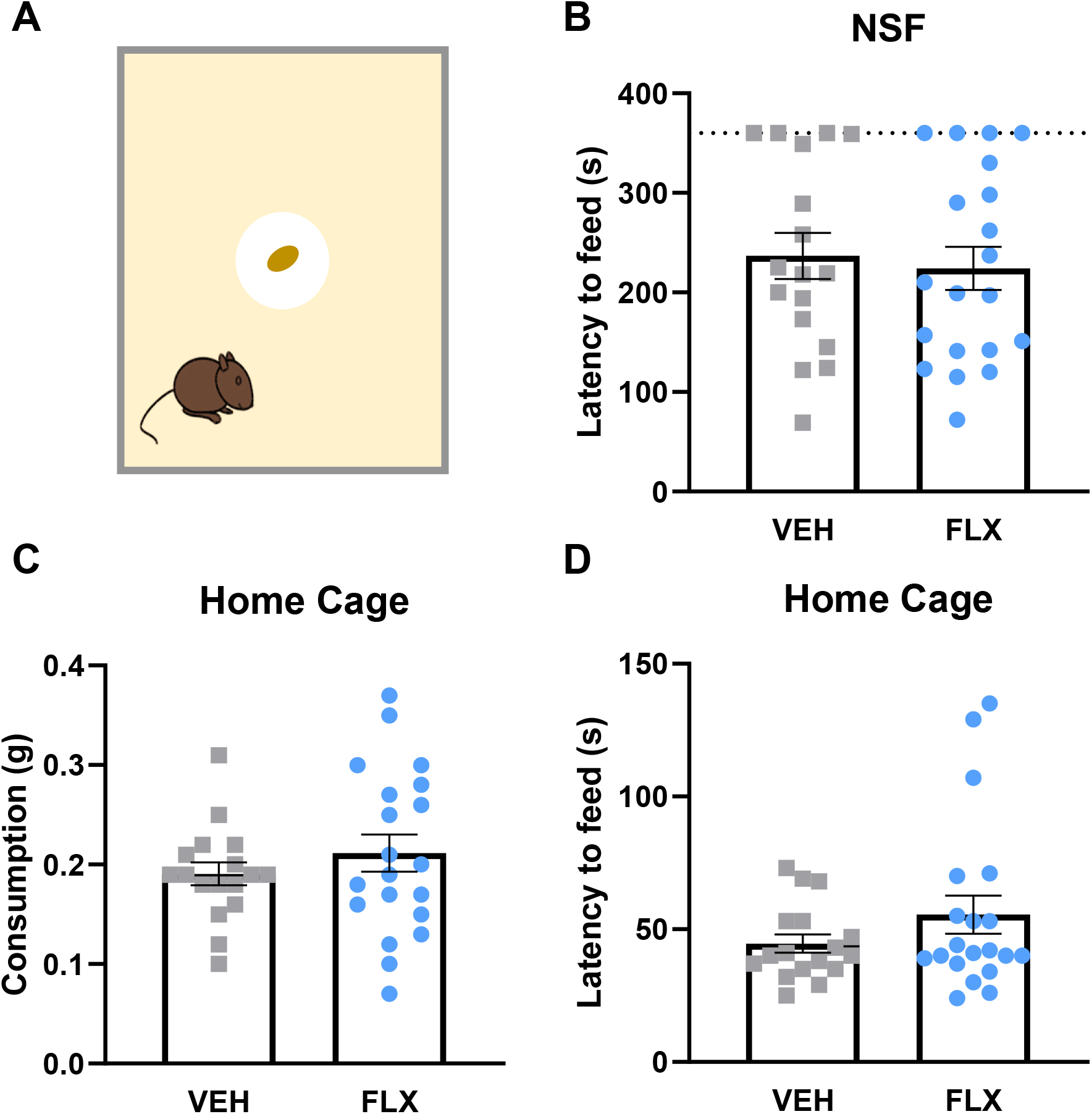
No effect on novelty suppressed feeding behavior following adult fluoxetine treatment in post-natal fluoxetine mice. **A.** Diagram of the behavior paradigm. **B.** Latency to feed in NSF. Dotted line at 360 seconds shows the behavior time out. Unpaired, two-tailed, t-test, p=0.6962. **C.** Amount of food consumed in the home cage in a 5-minute period. Unpaired, two-tailed, t-test, p=0.3650. **D.** Latency to eat in the home cage. Unpaired, two-tailed, t-test, p=0.2044. Gray squares represent adult vehicle-treated PN FLX animals, N=17. Blue circles represent adult fluoxetine-treated PN FLX animals, N=20. Bars represent mean. Error bars represent standard error.

To determine whether the paradoxical anxiogenic response to chronic FLX in PN-FLX mice was specific to C57 animals, we repeated these chronic adult FLX administration experiments in PN-FLX and PN-VEH mice on a 129 background, as well as in a cohort of naïve 129 mice. Similar to what was seen in PN-FLX C57 mice, chronic FLX administration of PN-FLX 129 mice in adulthood also increased avoidant behaviors. PN-FLX 129 mice treated chronically with FLX in adulthood showed an increased latency to feed in the NSF compared to their chronically VEH-treated counterparts (Fig. S4a; unpaired t-test, *P = 0.04, n = 27 VEH, 30 FLX, sexes combined). PN-VEH 129 mice treated chronically with FLX showed no difference in latency to feed compared to those treated chronically with VEH as adults (Fig. S4c; unpaired t-test, P = 0.88, n = 36 VEH, 32 FLX, sexes combined). In contrast, naïve 129 mice treated chronically with FLX as adults showed the classical anxiolytic response, with a decreased latency to feed compared to those chronically treated with the VEH (Fig. S4e; unpaired t-test, *P = 0.03, n = 9 VEH, 8 FLX, sexes combined). Again, these results were not due to differential effects of hunger as chronic FLX administration did not affect the amount of food consumed in the home cage by each group(Fig. S4b; unpaired t-test, P = 0.34, n = 27 VEH, 30 FLX, sexes combined; S4d; unpaired t-test, P = 0.95, n = 36 VEH, 32 FLX, sexes combined; S4f; unpaired t-test, P = 0.18, n = 9 VEH, 8 FLX, sexes combined).

Cumulatively, these results suggest that rather than being anxiolytic, chronic FLX administration in adulthood is anxiogenic in mice that received early postnatal FLX exposure.

### Chronic tianeptine decreases avoidant behavior in adult mice postnatally exposed to fluoxetine

Following a two-week washout period, we pseudorandomly assigned the C57 PN-FLX and PN-VEH mice into chronic TIA and VEH administration groups (Fig. 3a; shown in green). Mice were administered 30 mg/kg TIA intraperitoneally 2x per day for 14 days or a saline vehicle. This dose of TIA was chosen as it has previously been shown to normalize avoidant, motivated and hedonic behaviors in mice chronically treated with corticosterone^18^, and unpublished data from our lab has shown that 14 days of administration is sufficient to see these behavioral changes. Chronic TIA increased rearing in PN-FLX mice in both the first 10 minutes (Fig. 3d; unpaired t-test, *P = 0.02, n = 19 VEH, 20 TIA, sexes combined) as well as the entire 60 minutes (Fig. S5b; unpaired t-test, **P = 0.005, n = 19 VEH, 20 TIA, sexes combined). Chronic TIA also resulted in an increase in center time (Fig. 3e; unpaired t-test, *P = 0.03, n = 19 VEH, 20 TIA, sexes combined) in the first 10 minutes of the OFT. Chronic TIA also increased center time in PN-VEH mice in the open field (Fig. S6e,f). Chronic TIA administration had no effect on locomotor behavior in PN-FLX mice in the OFT (Fig. 3c; first 10 minutes, unpaired t-test, P = 0.77, n = 19 VEH, 20 TIA, sexes combined; Fig. S5a; total 60 minutes, unpaired t-test, P = 0.28, n = 19 VEH, 20 TIA, sexes combined). These findings suggest that chronic TIA administration does not grossly impact motor function but does increase novelty-induced exploration in PN-FLX mice, and decrease avoidance behavior broadly.

**Figure 3.**
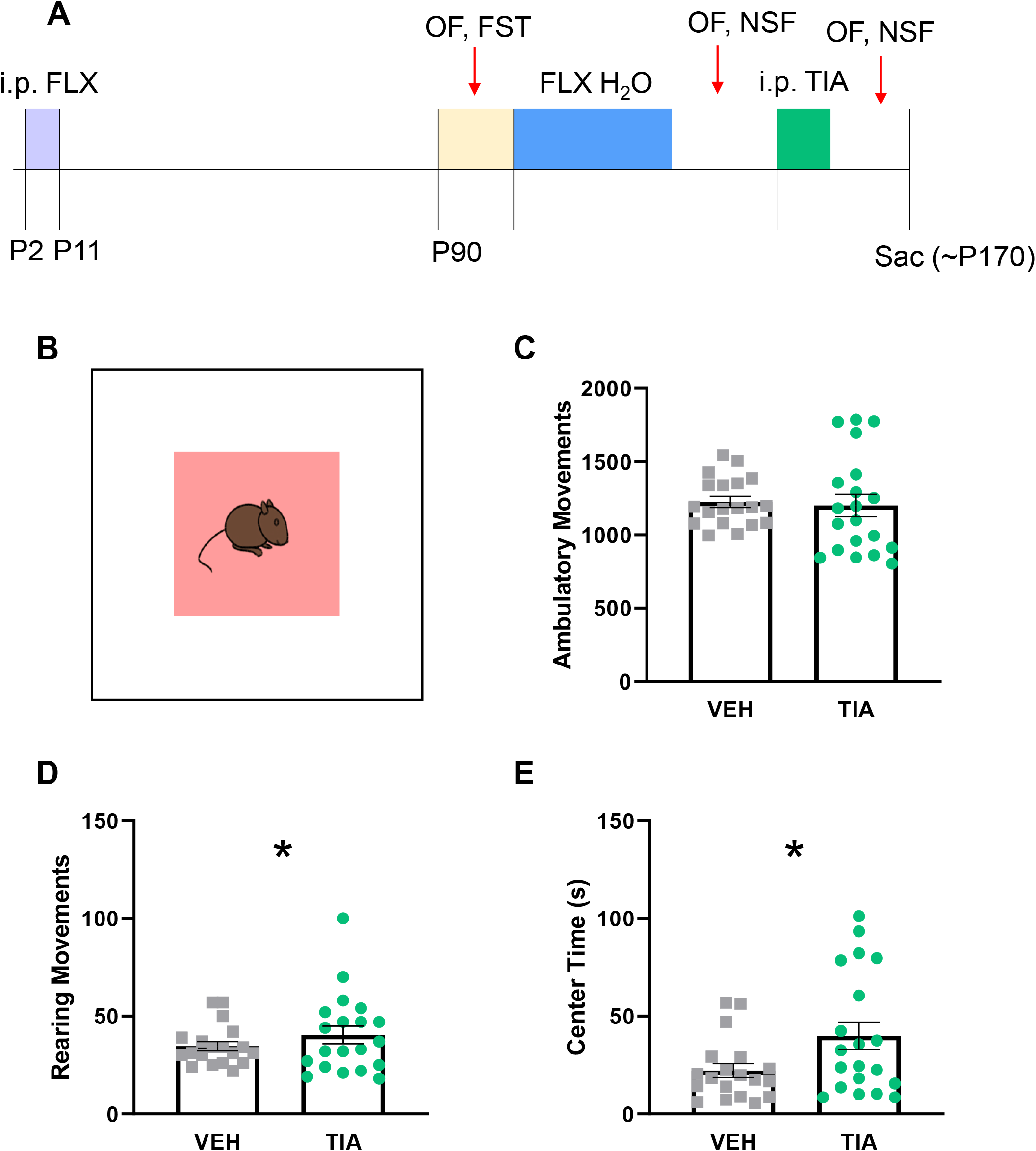
Decreased anxiety-like behavior in the first 10 minutes of open field following adult tianeptine treatment in post-natal fluoxetine mice. **A.** Timeline of experiment. Green shows the period during which tianeptine was administered after which follow up tests were performed. OF (open field), FST (forced swim test), NSF (novelty suppressed feeding). **B.** Diagram of open field enclosure, with center shown in red. **C.** Ambulatory movements in first 10 minutes of open field. Unpaired, two-tailed, t-test, p=0.7698. **D.** Same as C, except time rearing is shown, unpaired two-tailed, t-test, p=0.0224. **E.** Same as C, except time spent in center is shown, unpaired, two-tailed, t-test, p=0.0316. Gray squares represent adult vehicle-treated PN FLX animals, N=19. Green circles represent adult tianeptine-treated PN FLX animals, N=20. Bars represent mean. Error bars represent standard error. *p<0.05.

We also repeated the NSF task following chronic TIA administration to test whether TIA would improve avoidant behaviors in PN-FLX mice. Adult PN-FLX mice chronically treated with TIA demonstrated a decreased latency to feed compared to those treated with VEH (Fig. 4b; unpaired t-test, ***P=0.0001, n=17 VEH, 19 TIA, sexes combined). TIA had no effect on home cage food consumption or latency to feed in PN-FLX mice when the sexes were combined (Fig. 4c; unpaired t-test, P=0.11, n=17 VEH, 19 TIA, sexes combined; Fig. 4d; unpaired t-test, P=0.5817, n=17 VEH, 19 TIA, sexes combined), confirming that the decreased latency to feed was not due to a change in hunger. Chronic administration with TIA in PN-VEH mice did not alter their latency to feed (Fig. S7), suggesting that TIA is primarily effective in C57 mice with a baseline affective phenotype. Cumulatively, these data suggest that unlike chronic FLX, chronic TIA normalizes avoidant symptoms in adult PN-FLX mice.

**Figure 4.**
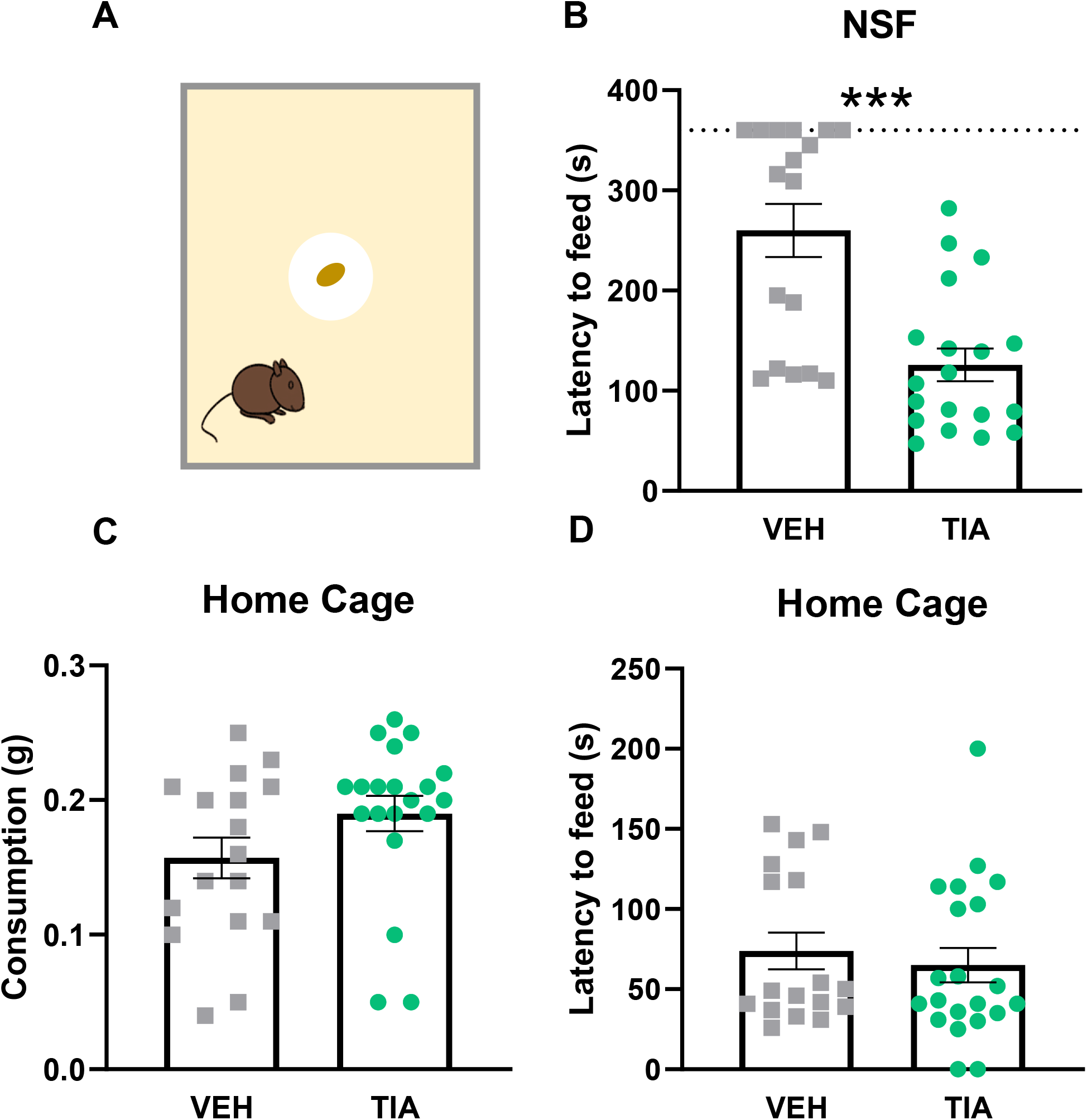
Decreased anxiety-like behavior in the novelty suppressed feeding task following adult tianeptine treatment in post-natal fluoxetine mice. **A.** Diagram of the behavior paradigm. **B.** Latency to feed in NSF. Dotted line at 360 seconds shows the behavior time out. Unpaired, two-tailed, t-test, p=0.0001. **C.** Amount of food consumed in the home cage in a 5-minute period. Unpaired, two-tailed t-test, p=0.1076. **D.** Latency to eat in the home cage. Unpaired, two-tailed, t-test, p=0.5817. Gray squares represent adult vehicle-treated PN FLX animals, N=17. Green circles represent adult fluoxetine-treated PN FLX animals, N=19. Bars represent mean. Error bars represent standard error. *p<0.05, ***p<0.001

Because of the crossover nature of these experiments, we verified that prior administration with chronic FLX or vehicle had no effect on the effect of TIA in OFT avoidance behaviors (rearing in first 10 min, three-way ANOVA, F(1, 61) = 3.06, P = 0.09, interaction of adult FLX and TIA administrations; rearing in 60 minutes, three-way ANOVA, F(1, 61) = 3.61, P = 0.06, interaction of adult FLX and TIA administrations; center time in first 10 minutes, three-way ANOVA, F(1, 61) = 0.70, P=0.41, interaction of adult FLX and TIA administrations; center time in 60 minutes, three-way ANOVA, F(1, 61) = 0.001, P = 0.98, interaction of adult FLX and TIA administrations) or the NSF (three-way ANOVA, F(1, 58) = 1.91, P = 0.17, interaction of adult FLX and TIA administrations) in PN-FLX mice.

## DISCUSSION

Our findings demonstrate the efficacy of TIA, an atypical antidepressant and MOR agonist, as an efficacious option in mice in which standard antidepressants that target the serotonin transporter, such as FLX, are unsuccessful. We found that early life exposure to FLX, leads to an increase in avoidant behaviors in male and female C57 mice, similar to what has previously been reported in 129 mice^9^ as well as in C57 mice treated for a more extended postnatal period^25^. Moreover, these affective symptoms in PN-FLX mice were not normalized by chronic administration with FLX in adulthood. Instead, chronic FLX exposure in adult PN-FLX mice exacerbated decreases in novelty-induced exploratory behavior, including decreased rearing behavior in the OFT, and increased avoidant behavior measured using center time in the first 10 minutes of this same test. In contrast, chronic administration with TIA in these same adult PN-FLX mice increased novelty-induced exploration and decreased avoidant behavior overall. These results suggest that TIA may be useful as a administration option for depression and anxiety in humans where direct targeting of the serotonergic system proves ineffective, and further, provide a framework for utilizing different etiologies of depression to direct the use of targeted therapies.

### Avoidant behavior following early fluoxetine exposure

While we found that PN-FLX C57 mice show decreases in novelty-induced exploration and increases in avoidance behavior, similar to effects seen in PN-FLX 129 mice, PN-FLX C57 mice did not show the same increases in behavioral despair seen in PN-FLX 129 animals. More pronounced behavioral effects in C57 mice in novelty-induced exploration and avoidant behavior, rather than anhedonic and despair-related behavior, are consistent with other developmental SSRI studies in mice. For example, C57 mice treated with FLX from P4-P22 showed reduced distance traveled and time spent moving in the OFT^25,31^. Similarly, C57 offspring gestationally exposed to FLX from E8-E18, showed an increased latency to feed in the NSF test as adults^32^. Moreover, these differences in the effects of PN-FLX on C57 and 129 mice are consistent with previous work demonstrating differing behavioral effects of deleting the serotonin transporter (5-HTT-KO) in mice of different genetic backgrounds. While 5-HTT-KOs on a C57 background showed more avoidant behavior, those on a 129 background showed more pronounced behavioral despair^33^. Cumulatively, comparing the effects of manipulations such as PN-FLX in different strains of mice contributes to our understanding of how genetic background can interact with environmental exposures to influence the presentation of behavioral symptoms associated with mood and anxiety disorders^8^.

### Adult fluoxetine administration increases avoidant behavior in PN-FLX mice

Intriguingly, chronic FLX administration did not improve, and even exacerbated, affective symptoms in PN-FLX mice. These changes included decreasing novelty-induced exploration, as indicated by ambulatory and rearing movements in the OFT, and increasing avoidant behaviors, measured as a decrease in center time in this same task. The paradoxical effects of chronic FLX administration on avoidant behavior in adult PN-FLX mice could occur because the serotonergic system appears persistently disrupted in these mice. For example, early postnatal fluoxetine exposure produces adult mice with a hyperactive median raphe, hypoactive dorsal raphe^11^ as well as altered prefrontal modulation of both serotonergic and non-serotonergic raphe neurons^10^. In fact, when serotonergic activity was pharmacologically inhibited in the median raphe in PN-FLX mice, avoidant behavior was decreased, implying that hyperactivity in the media raphe plays a key role in regulating this behavior in PN-FLX animals^11^. It is possible that adult SSRI administration is ineffective at relieving, and possibly aggravates, avoidant and anhedonic behaviors because of this different baseline set-point of serotonergic neurons in PN-FLX mice.

Of note, chronic FLX usually only ‘improves’ avoidant, motivated and despair-related behaviors in C57 mice whose emotional behavior is altered at baseline, such as by exposure to chronic corticosterone^21,23^. Therefore, it is possible that our C57 PN-FLX mice were simply not ‘stressed’ enough for chronic FLX administration to decrease avoidant behavior. However, even on a 129 background where PN-FLX mice have a very pronounced affective behavioral phenotype^9,34^, chronic adult FLX administration paradoxically increased avoidant, and decreased hedonic behavior. Considering naïve 129 mice show a decrease in avoidant behavior and an increase in hedonic behavior in response to chronic FLX, these findings in 129 PN-FLX mice indicate that postnatal FLX exposure creates an insensitivity to adult FLX administration regardless of the extent of baseline emotional behavioral changes.

Our findings differ from those of Karpova et al, who found that adult chronic FLX exposure improved several measures of avoidant behavior in mice postnatally treated with FLX from P4-P21^25^. Although in 129 mice P2-P11 and P4-P21 PN FLX administration produce many of the same adult behavioral differences, Rebello et al showed that in the elevated plus maze, the effects of P2-P11 and P12-P24 PN FLX are actually diametrically opposed, with P2-P11 being anxiogenic and P12-P24 being anxiolytic^9^. Thus, differences in the length of PN FLX exposure may explain the differences in our findings. As the P2-P11 window better approximates the brain development occurring during the third trimester of a human pregnancy, we believe our results would likely be more to the human condition.

### Adult tianeptine administration decreases avoidant behavior in PN-FLX mice

In contrast to chronic FLX, chronic TIA administration increased novelty-induced exploratory behavior and motivation in the NSF, and decreased avoidant behavior in both the OFT and NSF in adult PN-FLX mice. In the OFT, chronic TIA resulted in a significant increase in rearing and time spent in the center during the first 10 minutes in adult PN-FLX mice, without increasing overall ambulation. These behaviors cannot be attributed to a direct psychostimulant effect of tianeptine as the drug was administered over 14 hours prior to all behavioral tests. As tianeptine clears from both blood plasma and brain tissue within 2 hours^18^, the behavioral changes we observe must result from chronic administration as opposed to acute effects.

The ability of chronic TIA to decrease avoidant behaviors in PN-FLX mice, while chronic FLX does not, may be due to persistent alterations in their serotonergic system that make them differentially sensitive to the mechanisms targeted by TIA versus FLX. FLX’s proposed mechanisms of action include blocking serotonin reuptake, and antagonizing 5-HT1A autoreceptors, both of which increase monoamine levels over time. Increased monoamines, may in turn allow for enhanced plasticity and hippocampal neurogenesis, which also contribute to the drug’s ultimate behavioral efficacy^6^. By contrast, TIA does not directly target the monoaminergic system and instead has been shown to be a MOR agonist^17,35^. MORs are necessary for the improvement in affective symptoms elicited by chronic administration of TIA in mice chronically administered corticosterone^18^, while acute depletion of serotonin does not affect TIA’s efficacy (Rene Hen, personal communication). TIA does not inhibit biogenic amine transporters, but has been shown in indirectly modulate glutamatergic transmission, increase plasma dopamine and promote neuroplasticity^35,36^. These differences may explain the differential behavioral efficacy of these two drugs in PN-FLX mice.

### Limitations

We found that chronic adult administration with TIA decreases some measures of avoidant behavior in PN-VEH mice relative to adult vehicle-treated controls. This could indicate that TIA is capable of exerting an anxiolytic effect in individuals who do not display an anxiety- or depression-like phenotype at baseline. However, the efficacy of TIA in the control PN-VEH mice may also be due to residual effects of stress experienced following repeated chronic injections during early postnatal life.

This is emphasized when comparing naïve 129 mice versus PN-VEH 129 mice that were then treated with FLX in adulthood – FLX administration had no effect on latency to feed in NSF in PN-VEH 129s, but significantly decreased this time in naïve 129s (Fig. S3), indicating that post-natal injection stress may have some effect on its own in this model. This may be beneficial for investigations into mouse models of depression as human patients often develop affective disorders following multiple ‘hits’ of stress or adversity^37^.

Finally, in our experiment the same cohort of PN-FLX or PN-VEH animals first underwent chronic administration with FLX or its respective vehicle, followed by a subsequent chronic administration with either TIA or its respective vehicle administration. While it is possible that the first drug administration could have influenced the outcome of the second, we believe this is unlikely as we randomly reassigned the animals to each of the administration or vehicle conditions and allowed a two-week washout period between the chronic administrations. We did not observe an effect of crossover using direct statistical assessment. Moreover, this repeated administration paradigm – first with an SSRI followed by another drug - most closely matches the experience of treatment-resistant patients as current front-line treatments for depression are SSRIs, with alternate medications considered after SSRIs have failed.

### Broader implications

The goal of precision medicine is to identify the most effective treatment for a given patient based on known risk factors, biomarkers, or specific genes^38^. An example in the treatment of depression is genetic testing for *CYP2D6* and *CYP2C19* genes to help decide what dose of the antidepressant amitriptyline a patient might need, as these genes regulate liver enzymes that can lead to slow or fast metabolism of the drug^39^. By understanding how developmental serotonin disruption can affect adult antidepressant treatment response, we hope to productively inform treatment decisions for patients with a history of this early exposure.

Treatment resistance is a product of not knowing for whom certain treatments would be most effective. Here we show resistance to adult FLX administration in a developmental FLX exposure model. However, it remains unclear if early exposure to other SSRIs, such as escitalopram, paroxetine, and sertraline would also have these effects. A cohort of human patients was investigated for the effect of maternal SSRI use during pregnancy on the adult affective disorder rates of the children, however these rates are not broken down by specific drug treatment^12^ and there may be variability in these effects. This would be invaluable information for pregnant patients when making decisions about pharmacological treatment for depression. Further, this model can be expanded to examine possible developmental consequences of exposure to other monoaminergic drugs, for example norepinephrine– dopamine reuptake inhibitors, as well as adult treatment response. In addition to being clinically relevant, this would also expand our understanding of how monoamines impact development to affect adult mechanisms underlying behavior.

The efficacy of TIA at normalizing avoidant behavior in this PN-FLX model opens space for investigating alternative treatment targets to monoaminergic systems in the treatment of depression. Future experiments should examine non-monoamine antidepressants like tianeptine in other models of treatment resistance. Currently, ketamine is the only FDA approved non-monoaminergic treatment for depression in the US. While the approval of ketamine in 2019 increased the number of options available for patients, this treatment still has limitations, including a highly restricted administration system and intense side effects including sedation^40^. It is important to continue to look for options that target other systems, including the opioid system, which has long been implicated in regulating mood and emotional behavior.

In summary, here we show that tianeptine may be a promising alternative treatment from SSRIs for humans with suspected *in utero* early developmental exposure to SSRIs, and more broadly that it may be helpful in a subset of treatment resistant depression in which SSRIs have not been successful.

## FUNDING AND DISCLOSURE

This work was funded by a grant from the Hope for Depression Research Foundation to Dr. Javitch. Dr. Javitch is co-inventor on patents held by Columbia University on tianeptine analogs. Dr. Canetta, Dr. Gingerich, and Dr. Ansorge declare no potential conflict of interest.

## FIGURE LEGENDS

**Supplementary Figure 1. Full 60-minute open field session behavior following adult fluoxetine treatment in post-natal fluoxetine mice**. **A.** Ambulatory movements are summed for all 60 minutes of open field. Unpaired, two-tailed, t-test, p=0.7012. **B.** Same as A, except rearing movements are shown, unpaired, two-tailed, t-test, p=0.2847. **C.** Same as A, except time spent in center is shown, unpaired, two-tailed, t-test, p=0.0745. Gray squares represent adult vehicle-treated PN FLX animals, N=17. Blue circles represent adult fluoxetine-treated PN FLX animals, N=20. Bars represent mean. Error bars represent standard error.

**Supplementary Figure 2. Open field behavior following adult fluoxetine treatment in post-natal vehicle mice**. **A.** Ambulatory movements in first 10 minutes of open field. Unpaired, two-tailed, t-test, p=0.001. **B.** Same as A, but ambulatory movements are summed for all 60 minutes of open field. Unpaired, two-tailed, t-test, p=0.0014. **C-D.** Same as A and B, except rearing movements are shown. C, unpaired two-tailed, t-test, p=0.0188. D, unpaired, two-tailed, t-test, p=0.0103. **E-F.** Same as A and B, except time spent in center is shown. E, unpaired, two-tailed, t-test, p=0.5455. F, unpaired, two-tailed, t-test, p=0.0418. Gray squares represent adult vehicle-treated PN VEH animals, N=15. Orange circles represent adult fluoxetine-treated PN VEH animals, N=16. Bars represent mean. Error bars represent standard error. *p<0.05, **p<0.01.

**Supplementary Figure 3. Novelty suppressed feeding behavior following adult fluoxetine treatment in post-natal vehicle mice and 129 mice**. **A.** Diagram of the behavior paradigm. **B.** Latency to feed in NSF. Dotted line at 360 seconds shows the behavior time out. Unpaired, two-tailed, t-test, p=0.1310. **C.** Amount of food consumed in the home cage in a 5-minute period. Unpaired, two-tailed, t-test, p=0.0366. **D.** Latency to eat in the home cage. Unpaired, two-tailed, t-test, p=0.2587. Gray squares represent adult vehicle-treated PN VEH animals, N=15. Orange circles represent adult fluoxetine-treated PN VEH animals, N=16. Bars represent mean. Error bars represent standard error. *p<0.05.

**Supplementary Figure 4. Novelty suppressed feeding behavior following adult fluoxetine treatment in post-natal fluoxetine, post-natal vehicle, and post-natal treatment naïve 129 mice**. **A.** Latency to feed in NSF in PN FLX mice. Dotted line at 600 seconds shows the behavior time out. Unpaired, two-tailed, t-test, p=0.0381. **B.** Amount of food consumed in the home cage in a 5-minute period in PN FLX mice. Unpaired, two-tailed, t-test, p=0.3405. Gray squares represent adult vehicle-treated PN FLX animals, N=27. Blue circles represent adult fluoxetine-treated PN FLX animals, N=30. **C-D.** Same as A-B, but in PN VEH animals. Dark gray squares represent adult-vehicle PN VEH animals, N=36. Orange circles represent adult fluoxetine-treated PN VEH animals, N=32. C. Unpaired, two-tailed, t-test, p=0.8830. D. Unpaired, two-tailed, t-test, p=0.9524. **E-F.** Same as A-B, but in PN naïve animals. Black gray squares represent adult vehicle-treated PN naïve animals, N=9. Teal circles represent adult fluoxetine-treated PN naïve animals, N=8. E. Unpaired, two-tailed, t-test, p=0.0324. F. Unpaired, two-tailed, t-test, p=0.1772. Bars represent mean. Error bars represent standard error. *p<0.05.

**Supplementary Figure 5. Decreased anxiety-like behavior in the full 60-minute open field session following adult tianeptine treatment in post-natal fluoxetine mice**. **A.** Ambulatory movements summed for all 60 minutes of open field. Unpaired, two-tailed, t-test, p=0.2774. **B.** Same as A, except time rearing is shown, unpaired, two-tailed, t-test, p=0.0053. **C.** Same as A, except time spent in center is shown, unpaired, two-tailed, t-test, p=0.0002. Gray squares represent adult vehicle-treated PN FLX animals, N=19. Green circles represent adult tianeptine-treated PN FLX animals, N=20. Bars represent mean. Error bars represent standard error. **p<0.01, ***p<0.001.

**Supplementary Figure 6. Open field behavior following adult tianeptine treatment in post-natal vehicle mice**. **A.** Ambulatory movements in first 10 minutes of open field. Unpaired, two-tailed, t-test, p=0.1207. **B.** Same as A, but ambulatory movements are summed for all 60 minutes of open field. Unpaired, two-tailed, t-test, p=0.0678. **C-D.** Same as A and B, except rearing movements are shown. C, unpaired two-tailed, t-test, p=0.0295. D, unpaired, two-tailed, t-test, p=0.0706. **E-F.** Same as A and B, except time spent in center is shown. E, unpaired, two-tailed, t-test, p=0.0084. F, unpaired, two-tailed, t-test, p=0.0188. Gray squares represent adult vehicle-treated PN VEH animals, N=14. Purple circles represent adult tianeptine-treated PN VEH animals, N=16. Bars represent mean. Error bars represent standard error. *p<0.05.

**Supplementary Figure 7. Novelty suppressed feeding behavior following adult tianeptine treatment in post-natal vehicle mice**. **A.** Diagram of the behavior paradigm. **B.** Latency to feed in NSF. Dotted line at 360 seconds shows the behavior time out. Unpaired, two-tailed, t-test, p=0.9829. **C.** Amount of food consumed in the home cage in a 5-minute period. Unpaired, two-tailed, t-test, p=0.2540. **D.** Latency to eat in the home cage. Unpaired, two-tailed, t-test, p=0.8779. Gray squares represent adult vehicle-treated PN VEH animals, N=15. Purple circles represent adult tianeptine-treated PN VEH animals, N=15. Bars represent mean. Error bars represent standard error.

## Notes

### Competing Interest Statement

The authors have declared no competing interest.

