## Supplementary figures and images for "Tianeptine, but not fluoxetine, decreases avoidant behavior in a mouse model of early developmental exposure to fluoxetine"

### Supplemental Figures

#### Slide 1
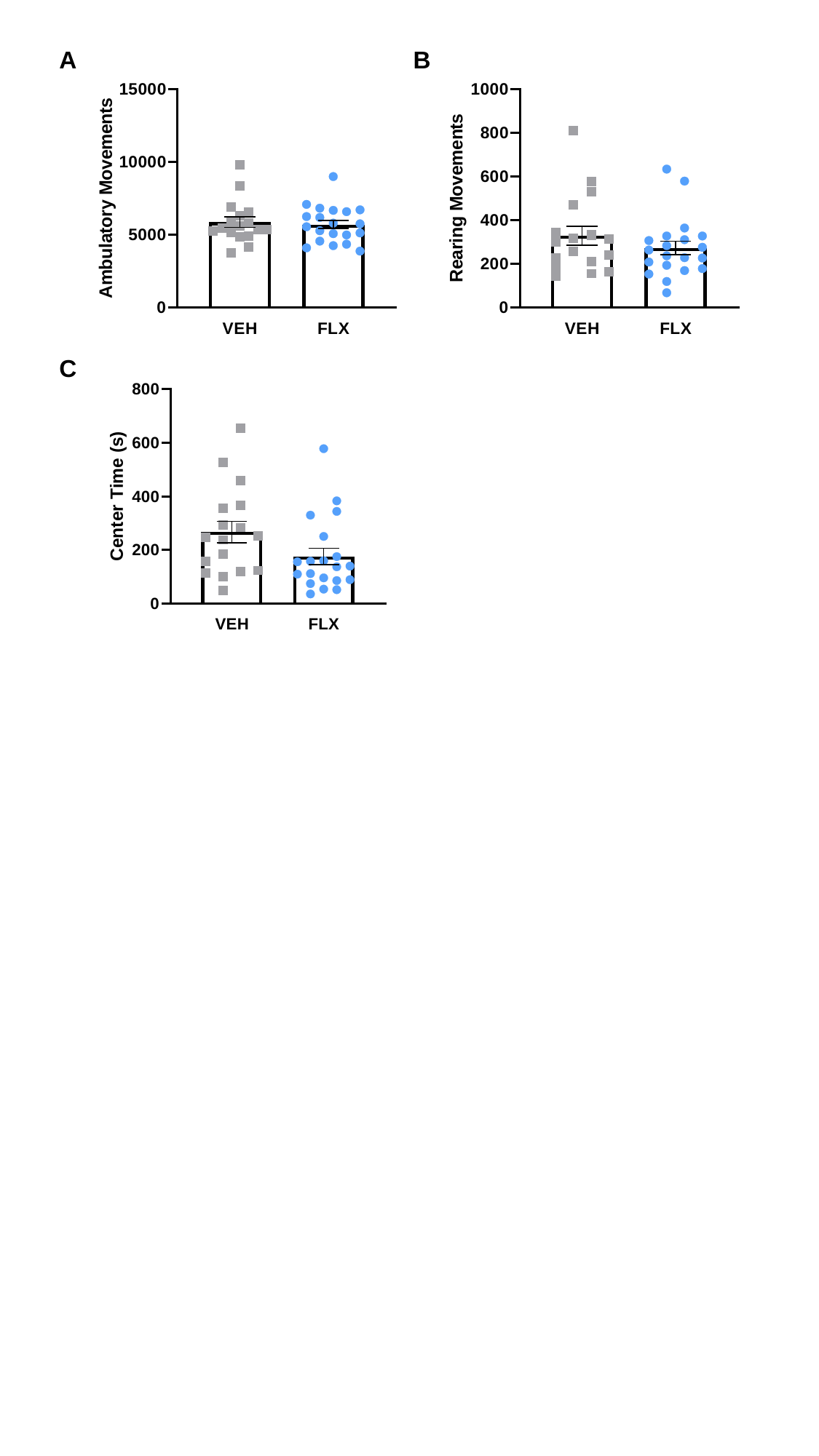

A
B
C

#### Slide 2
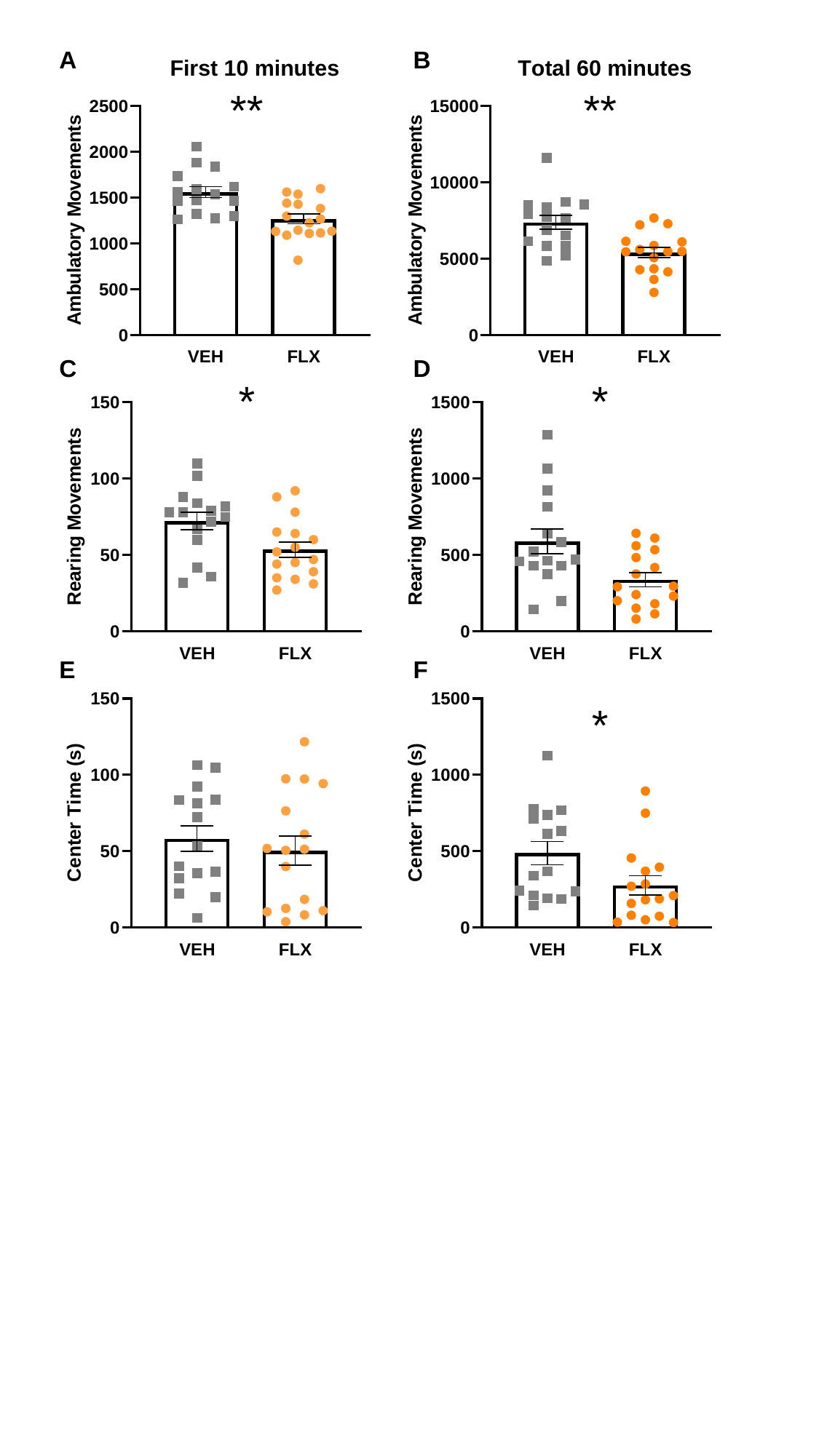

A
B
**
**
C
D
*
*
E
F
*

#### Slide 3
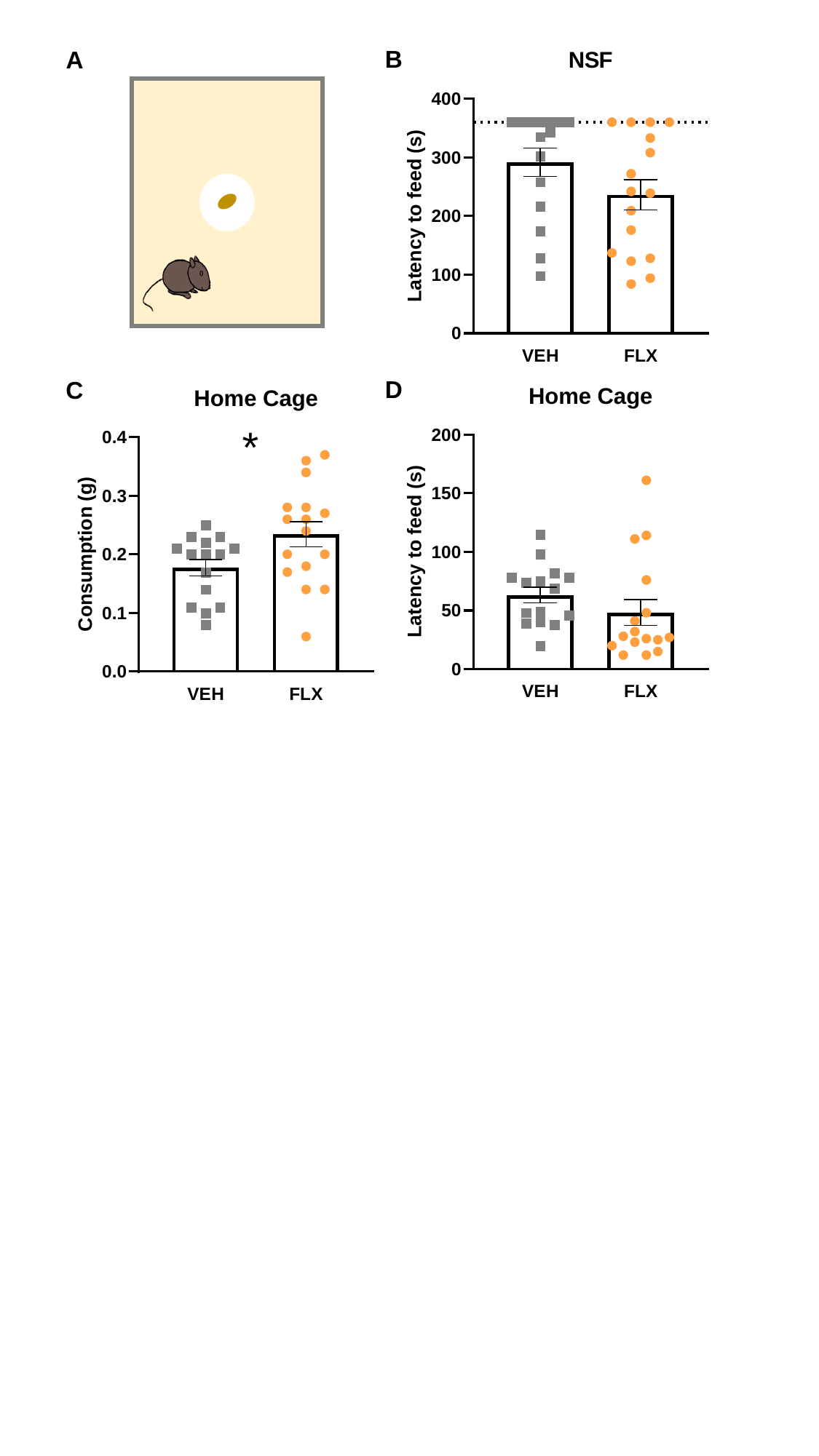

B
A
D
C
*

#### Slide 4
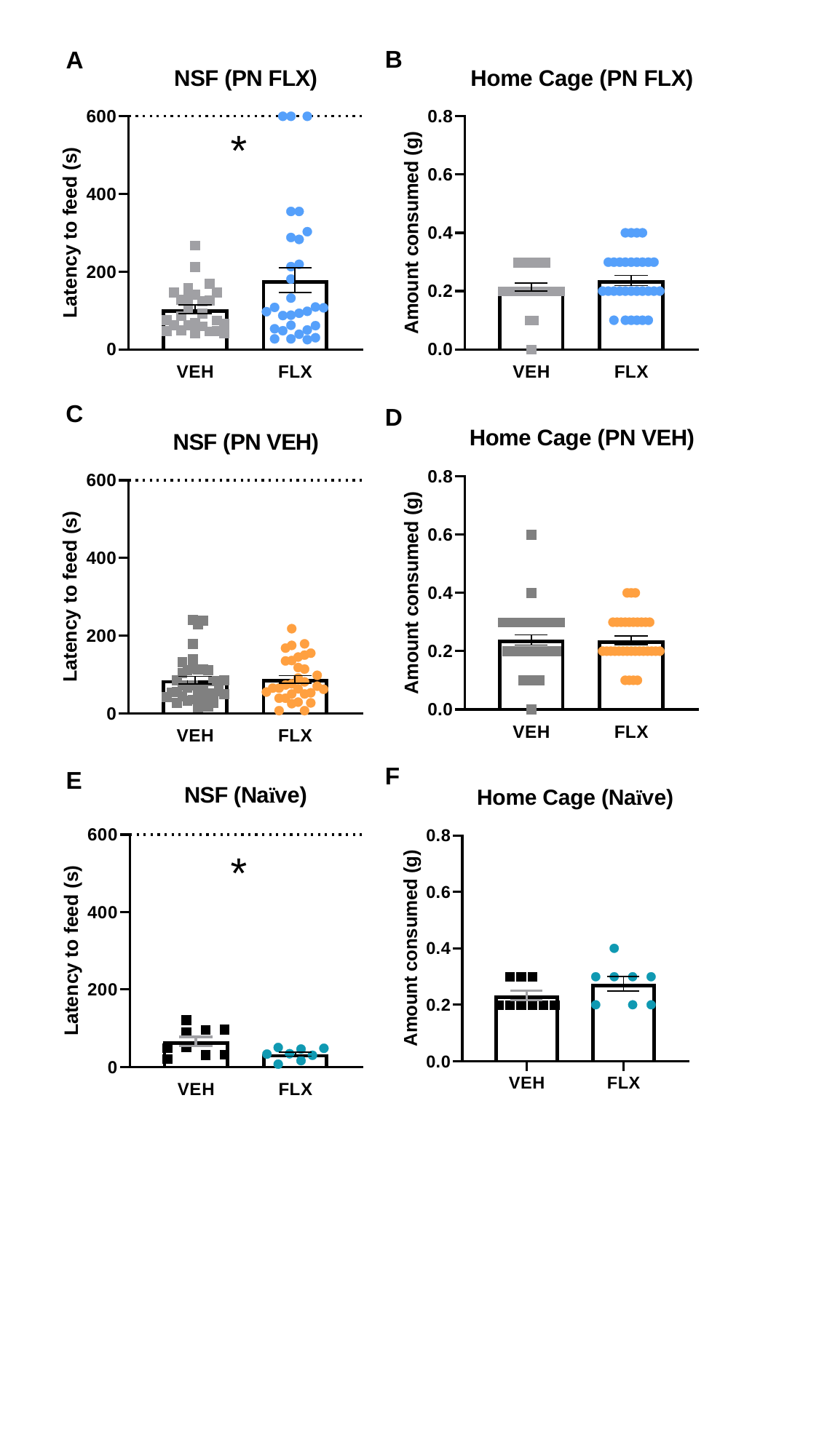

B
A
*
C
D
F
E
*

#### Slide 5
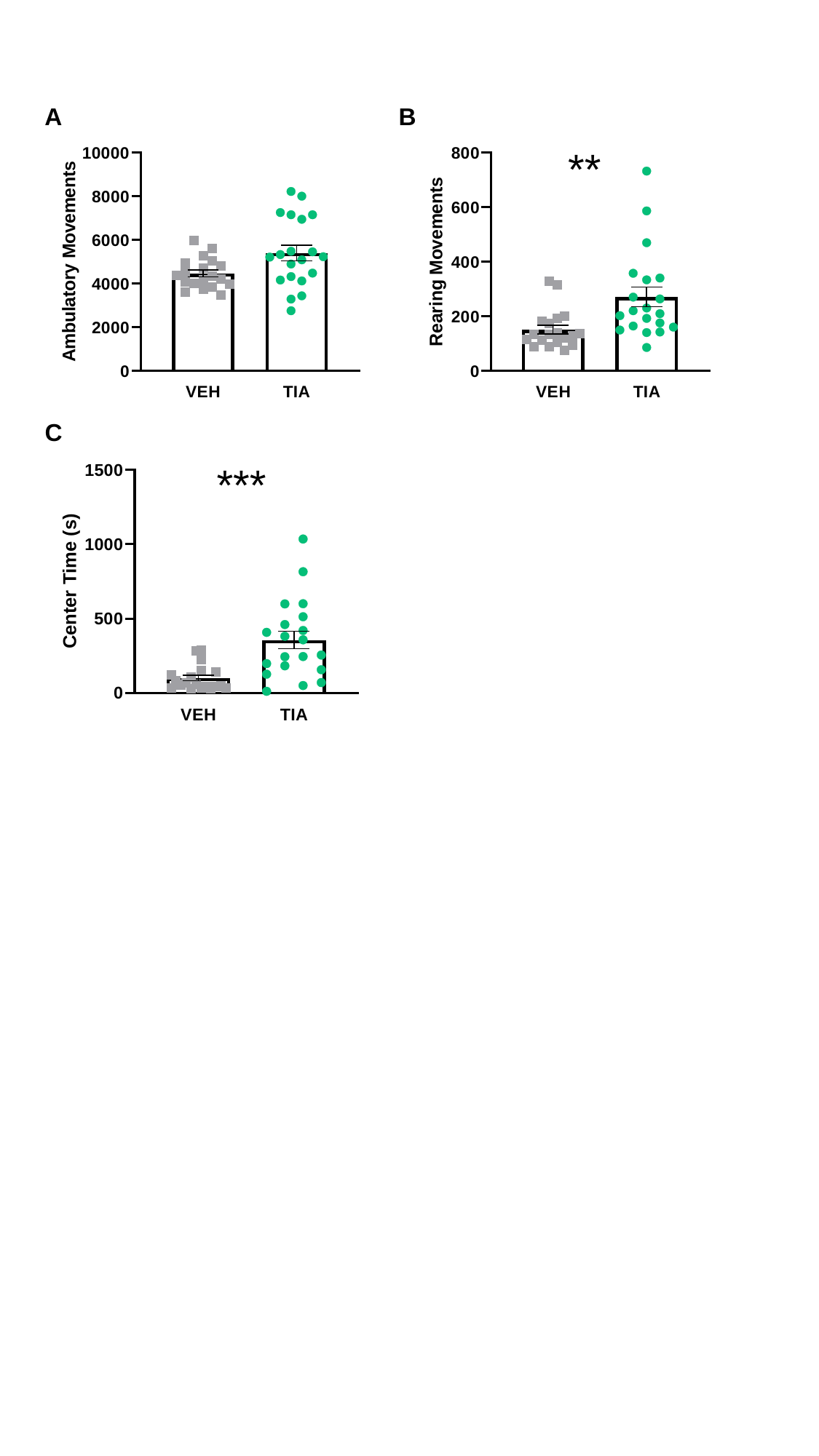

A
B
**
C
***

#### Slide 6
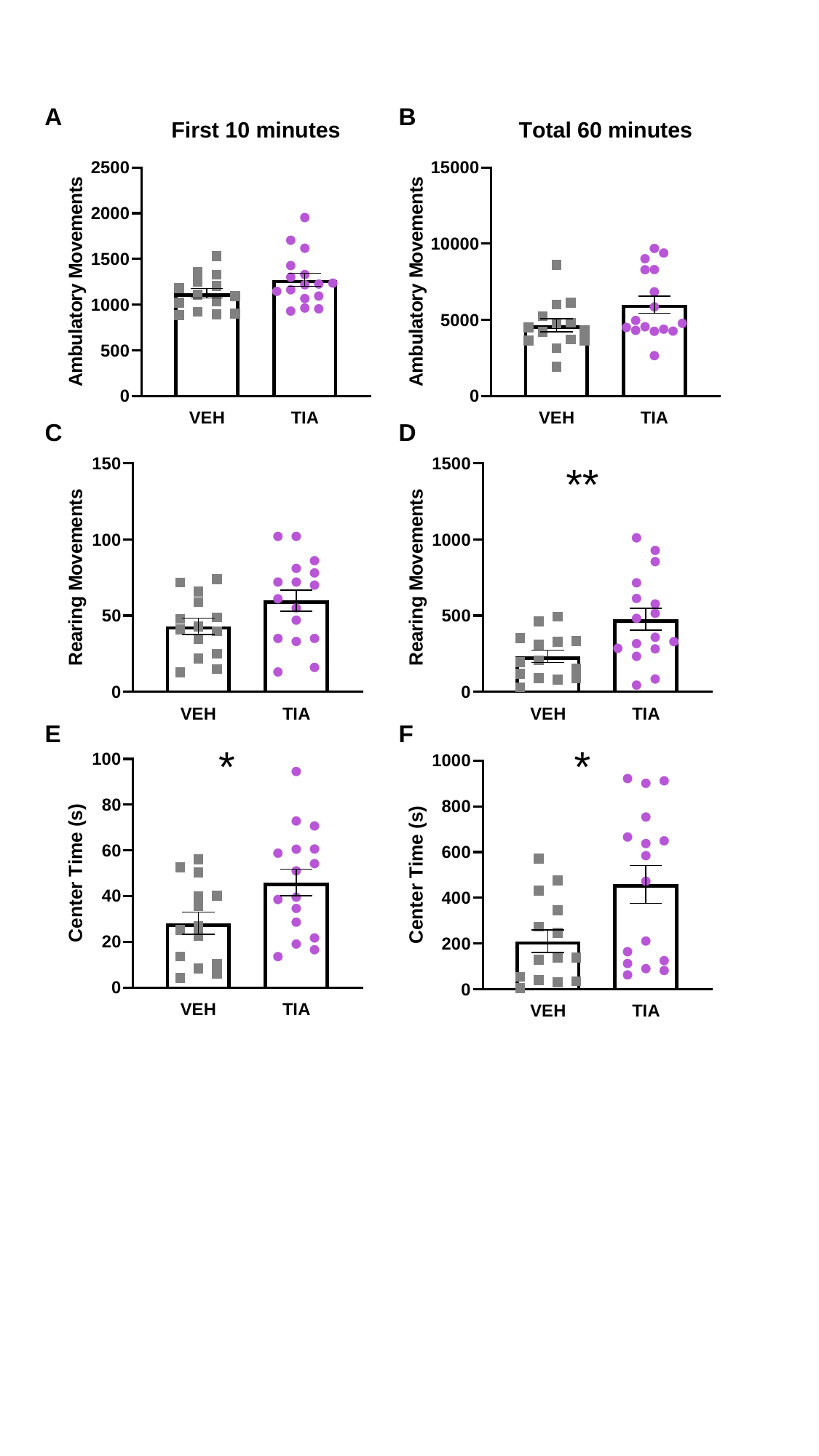

A
B
C
D
E
F
**
*
*

#### Slide 7
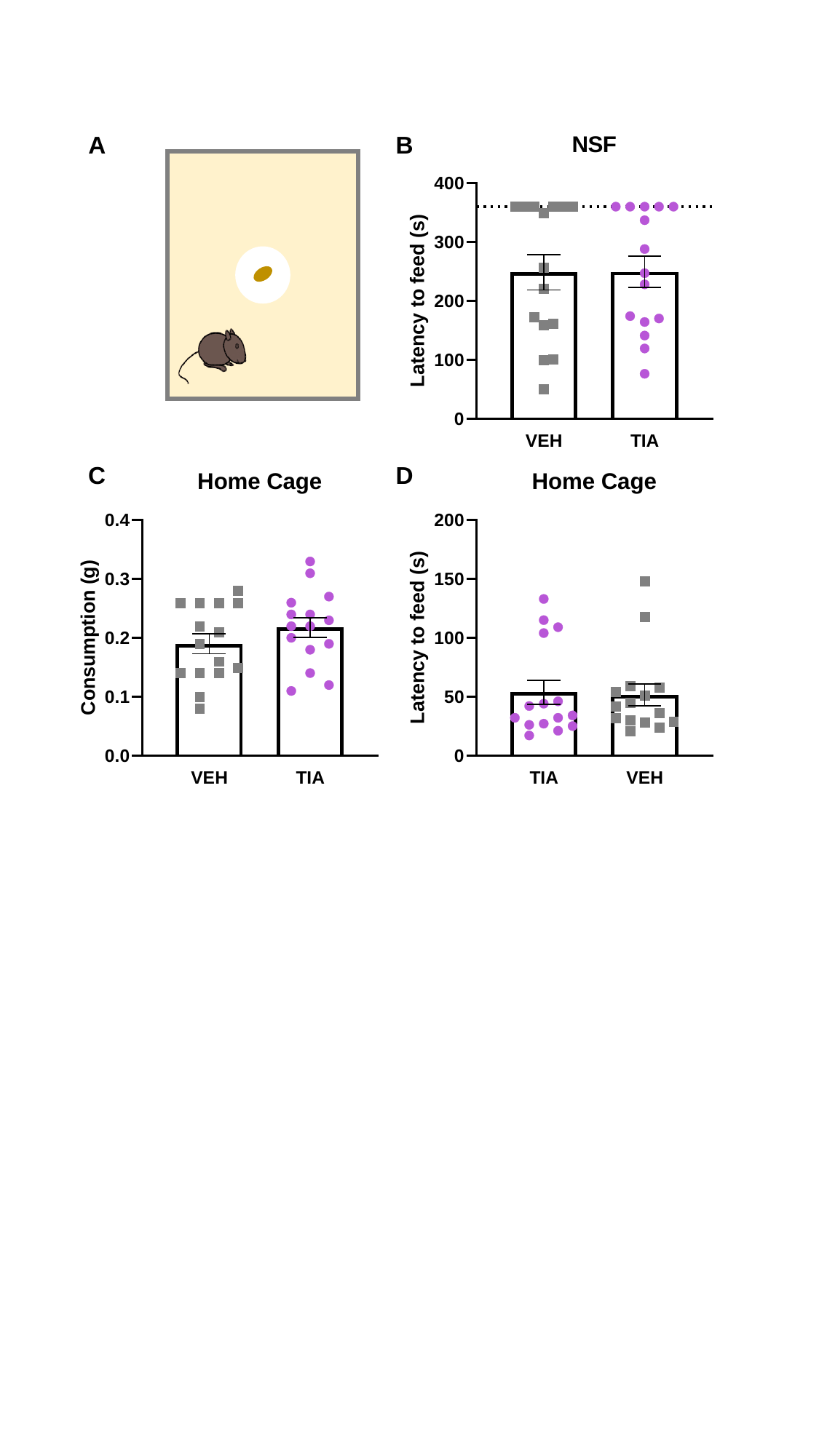

A
B
C
D
